# Behavioral and Brain Responses to Language Reflect Different Levels of Linguistic Representation

**DOI:** 10.64898/2026.08.21.746238

**Authors:** Andrea Gregor de Varda, Yevgeni Berzak, Evelina Fedorenko, Roger Levy

## Abstract

Human language processing can be studied through both behavior and brain activity, yet it remains unclear whether these two data types reflect sensitivity to the same information. One influential view holds that both behavioral and neural responses are largely determined by processing effort, often estimated by word surprisal together with the context-independent properties of word frequency and length. At the same time, neural responses have been shown to encode richer aspects of linguistic content, including meaning. Here, we use neural network language models to operationalize these alternatives and systematically compare, within the same analytic computational framework, the predictive power of low-dimensional effort-based predictors and high-dimensional embedding representations that encode contextualized linguistic content, including meaning. Across 8 behavioral datasets and 5 neural datasets (4 fMRI and 1 ERP), we find that processing effort captures substantial variance in both behavioral and neural measures of language processing, in line with much previous work. However, for brain responses—but not for behavioral measures—embedding representations carry substantial predictive power beyond the estimates of processing effort. These results therefore suggest that neural data provide access to rich, high-dimensional dynamics of language comprehension, whereas behavioral data reflect a bottlenecking of these dynamics into a small set of theoretically motivated properties of contextualized linguistic input.

**Significance Statement:** Two research communities study language comprehension as it unfolds in real time: psycholinguists use behavioral measures, such as eye movements during reading, and neuroscientists measure brain activity. The two are rarely studied together, but evidence from both must be integrated into a unified theory of language processing. Here we analyze both brain and behavioral responses within a single framework based on language models, comparing two long-standing accounts of what drives responses to language: processing effort versus meaning and other features not reducible to effort. We find that behavior is dominated by effort, whereas brain responses also reflect meaning. Developing a unified theory requires both kinds of data, but with a clear understanding of which levels of representation each measure reflects.

## 1 Introduction

Behavioral and brain responses both provide critical evidence for evaluating hypotheses about human language processing. However, although both are used to infer the latent computational mechanisms that underlie comprehension, they have largely been examined separately, by different scientific communities, and within different theoretical frameworks and methodological traditions. As a result, it remains a key open question what information each type of data encodes: do behavioral and neural responses reflect sensitivity to the same or distinct aspects of linguistic input? Answering this question is essential for

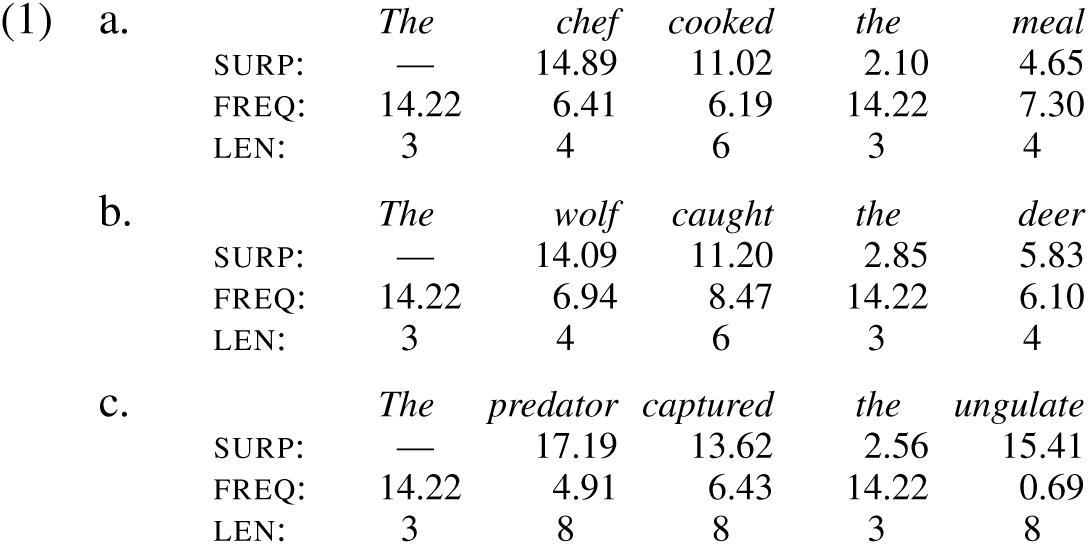

understanding how evidence from brain and behavior should be interpreted, compared, and ultimately integrated within a unified theory of language processing.

According to one view, both behavioral and neural responses to language are shaped by cognitive effort, and this effort is well captured by a small set of word-level predictors: word surprisal or contextual predictability (measured as negative log probability), word frequency, and word length (henceforth “surprisal, frequency, length”, SFL). These variables are central to longstanding theoretical accounts of processing cost in incremental language comprehension (Just and Carpenter, 1980; Rayner et al., 1996; Rayner and Duffy, 1986; Rayner, 1998; Clifton Jr et al., 2016). Indeed, cognitive effort, as operationalized through SFL, explains substantial variance in behavioral measures of language processing, particularly reading behavior (e.g., Schilling et al., 1998; Demberg and Keller, 2008; Smith and Levy, 2013; Berzak and Levy, 2023; Shain et al., 2024). Processing effort also accounts for a large amount of variance in brain responses, as seen in increased BOLD and ERP responses for more surprising or infrequent words during incremental comprehension or for sentences comprised of such words (Federmeier and Kutas, 1999; Frank et al., 2015; DeLong et al., 2005; Federmeier et al., 2007; Henderson et al., 2016; Brennan et al., 2020; Shain et al., 2020; Schmitt et al., 2021; Heilbron et al., 2022; Russo et al., 2022; Michaelov and Bergen, 2022; Michaelov et al., 2023; Tuckute et al., 2024b, among others). Finally, behavioral measures of comprehension difficulty during reading predict activity in language-responsive brain regions (Wehbe et al., 2021), suggesting that processing effort explains shared variance across brain and behavior.

However, effort alone cannot fully explain the processes underpinning human language processing. The primary function of the language system is to support the construction and interpretation of meaning, and traditional effort-based predictors abstract away from much of this content. Two sentences may have nearly identical profiles of surprisal, frequency, and length—and would therefore be predicted to elicit comparable processing effort—yet differ completely in meaning, as in “The chef cooked the meal” and “The wolf caught the deer” (cf. 1a and 1b). Conversely, two sentences may express similar meanings but differ substantially in these predictors, as in “The wolf caught the deer” versus “The predator captured the ungulate” (cf. 1b and 1c). Consistent with this intuition, a growing body of work shows that brain responses to language are modulated by semantic factors that cannot be straightforwardly reduced to processing effort, with perceptual (e.g., concreteness, imageability) and affective (valence, arousal) dimensions of meaning emerging as key organizing principles of neural representations (Huth et al., 2016; Deniz et al., 2019; Tuckute et al., 2024b, 2025). Behavioral measures of online language processing also show some degree of sensitivity to semantic properties (Juhasz and Rayner, 2003; Arfé et al., 2023; Yao et al., 2024), although these effects are typically weaker and less consistent than those observed in neural data (Magnabosco and Hauk, 2024). Together, these findings suggest that although effort-related signals are prominent in both kinds of data, brain responses may afford access to richer aspects of linguistic representation than behavior.

To test this possibility—that brain responses capture richer linguistic representations than behavior— we need a way to operationalize both i) effort-related properties and ii) representational features that encompass the meaning of words in context. Neural network language models (LMs) allow for such operationalization within a single computational framework. First, LMs can be used to derive next-word surprisal estimates—a core component of effort-based accounts of comprehension, together with the context-invariant properties of word frequency and length (our SFL predictors; Figure 1A). Such LM-derived quantities have been widely used to explain patterns in language processing behavior (Demberg and Keller, 2008; Goodkind and Bicknell, 2018; Frank and Hoeks, 2019; Wilcox et al., 2020; Kuribayashi et al., 2021; Merkx and Frank, 2021; Oh and Schuler, 2022; Berzak and Levy, 2023; Boyce et al., 2023; Shain et al., 2024; de Varda and Marelli, 2024). And second, these models generate high-dimensional internal representations—embeddings—that encode a wide range of linguistic properties, including form-and meaning-related information (Jawahar et al., 2019) that need not be reducible to surprisal, frequency, and length. Unlike SFL, these representations are not constrained by theories of effort; instead, they encode a host of linguistic properties learned to support prediction. Embedding representations (henceforth EMB) have been extensively used to account for patterns of brain activity during language comprehension (Jain and Huth, 2018; Gauthier and Ivanova, 2018; Gauthier and Levy, 2019; Sun et al., 2019; Schrimpf et al., 2021; Aw and Toneva, 2022; Oota et al., 2022; Caucheteux and King, 2022; Goldstein et al., 2022; Antonello et al., 2023; Tang et al., 2023; Tuckute et al., 2024b; de Varda et al., 2025; see Arana et al., 2023; Tuckute et al., 2024a for reviews; Figure 1B), but have rarely been applied to the prediction of linguistic behavior (but see Schrimpf et al., 2021; Hollenstein et al., 2022; Lin et al., 2025; Tsipidi et al., 2026). Further, past studies that have shown that model embeddings predict neural or behavioral language responses have not asked how much of the predictivity is due to the fact that the embeddings capture effort-relevant features (cf. Tuckute et al., 2024b).

**Figure 1:**
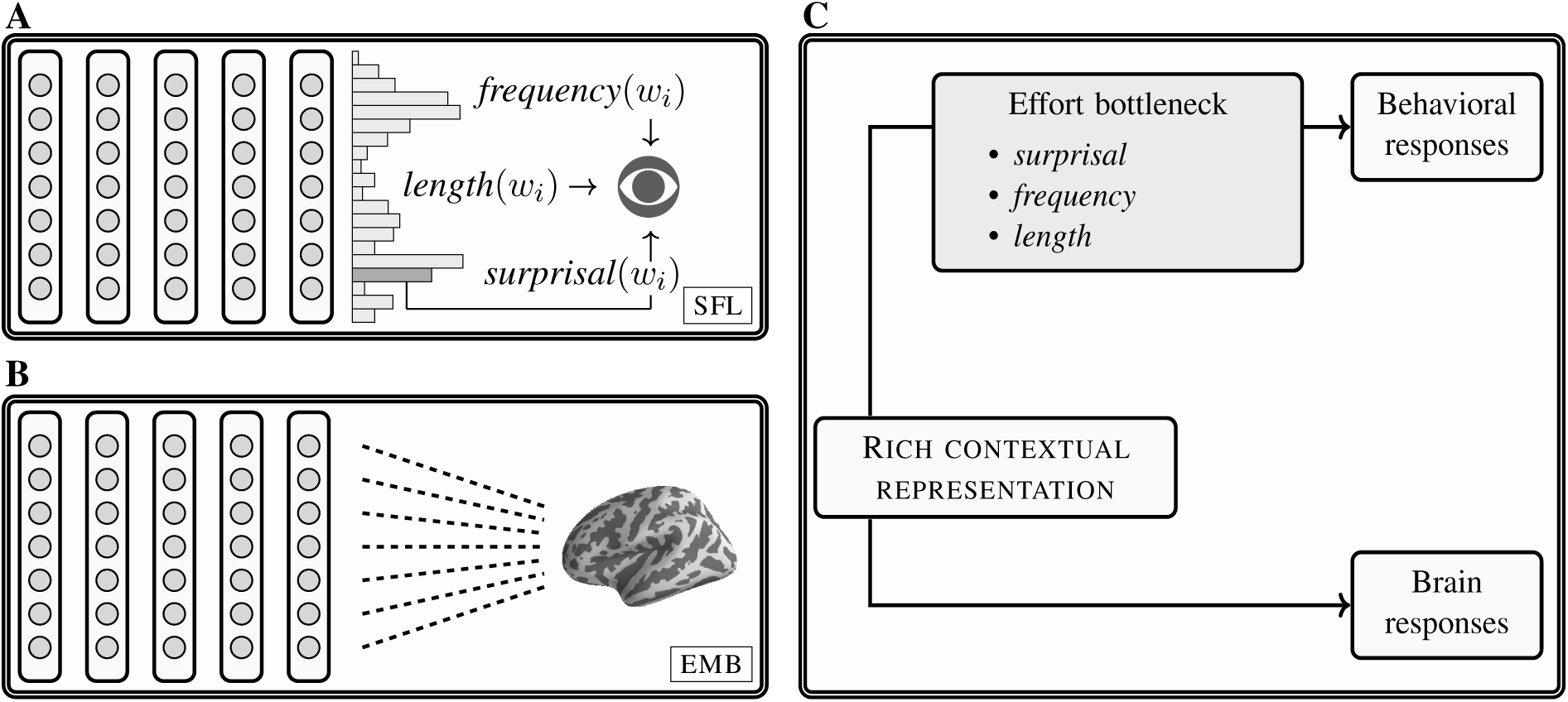
A, SFL: The model’s next-word probability yields *surprisal*, which together with word *frequency* and *length* is used to predict behavior. **B, EMB:** The model’s high-dimensional internal representations (embeddings) are directly used to predict brain responses. **C:** Hypothesized causal structure: contextual representations influence behavioral responses only through an effort bottleneck (*surprisal*, *frequency*, *length*), but they influence brain responses directly.

Here, we systematically investigate what kinds of information behavioral and neural measures of language processing are sensitive to. We hypothesize that behavioral responses reflect a *bottlenecking* process, whereby language representations are compressed into a small set of effort-related dimensions—surprisal, frequency, and length—which influence observable behavior (Levy, 2008). Under this proposal, high-dimensional representations shape behavior only indirectly, via their contribution to SFL. Conversely, we expect neural responses to reflect both effort-related signals captured by SFL and additional representational content captured by EMB but not by SFL, indicating more direct access to high-dimensional linguistic representations (Figure 1C). We test this hypothesis by comparing the predictive power of SFL, EMB, and their combination across a broad range of behavioral and neural datasets.

## 2 Results

We systematically compared the SFL and EMB approaches across behavioral and brain data. To this end, we used linear regression to predict human responses during sentence comprehension—as recorded using behavioral and neural measures—from either or both the SFL and EMB predictors:

1. SFL: surprisal, log-frequency, and length for the word *w*_i_.
2. EMB: the contextual representation of the word *w*_i_, i.e., the model’s activation in its final layer in response to *w*_i_ (we chose the final layer to capture comparatively high-level linguistic information including meaning; see Appendix A.5 for analyses considering all layers).
3. SFL^⌢^EMB: the concatenation of the two.

Our analyses were based on pre-existing datasets of language processing, including four eye-tracking datasets (Provo, MECO-En, ZuCo-2, UCL_ET_), three self-paced reading datasets (UCL_SPR_, Brown, NatStor_SPR_), one Maze (Forster et al., 2009; Boyce et al., 2020) dataset (NatStor_Maze_), one ERP dataset (UCL_N400_), and four fMRI datasets (Wehbe2014, NatStor_fMRI_, Pereira2018, Tuckute2024; see Figure 2 for an overview and Table 1 for details). For the majority of the datasets, responses were measured incrementally while participants read or listened to linguistic materials; for Pereira2018 and Tuckute2024, responses were recorded for sentences presented all at once, so a single measure per sentence was obtained. All brain and behavioral data were modeled as univariate responses, to ensure comparability across measures. For the behavioral datasets, response times were averaged across participants. For the fMRI data, we focused on a set of brain regions implicated in language processing (Hagoort, 2019; Fedorenko et al., 2024); within each region, we averaged BOLD responses across voxels, and then across regions and across participants. Finally, for the ERP data, we averaged the N400 amplitudes across 12 centro-parietal electrodes and across participants. Thus, for all behavioral and neural measures, we were left with a single value (average response) per word or sentence. We assessed the internal consistency of the measurements in each dataset using split-half reliability, which varied considerably across measures (mean = 0.51, range = 0.16–0.93; see Table 1 and Appendix A.7) due to differences in the number of participants and the noise properties of the different recording modalities.

**Figure 2:**
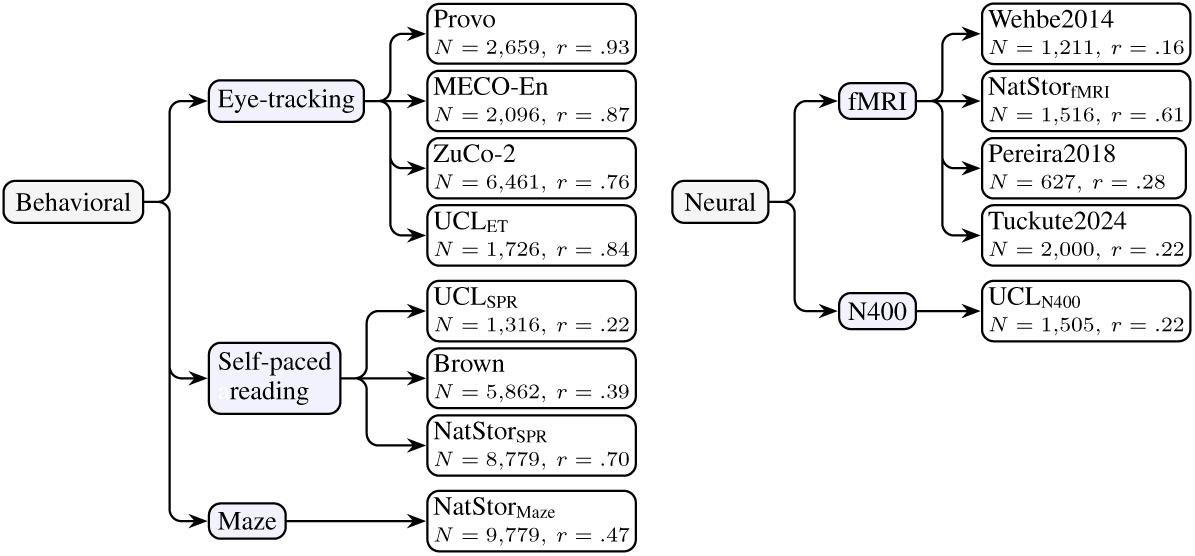
Overview of the behavioral (left) and neural datasets (right) used in the study. The arrows group the individual corpora under each measurement type. The values shown for each dataset are the number of observations included in the analyses (*N*, which is the number of words or sentences) and their split-half reliability (*r*, which is computed across participants).

**Table 1:** Summary of behavioral and neural datasets considered in the analyses. The columns *Type*, *Measure*, *Name*, and *Citation* report basic information to identify the datasets. The column *Resolution* indicates the granularity (words, sentences, TR intervals) of the responses. All the datasets are naturalistic, but some were edited to include specific constructions or cover certain topics; this information is reported in the column *Edited*. The datasets varied in the amount of connected text that was presented to models and participants, from sentences to a whole chapter; this information is specified in *Connected text*. *Modality* indicates the presentation modality; in the case of visual presentation, all connected text (sentences, passages, stories) was presented simultaneously unless stated otherwise. *N* refers to the actual number of data points included in the analyses, after data aggregation and trimming (see §5.3). *Part.* indicates the number of participants per data point (i.e., how many observations were used to calculate the mean response at the granularity level specified in *Resolution*). Split-half reliability is reported in the last column. See Appendix A.7 for details on how reliability varies across different sample sizes.

| Type | Measure | Name | Citation | Resolution | Edited | Connected text | Modality | N | Part. | Reliability |
| --- | --- | --- | --- | --- | --- | --- | --- | --- | --- | --- |
| Beh | Eye-tracking | Provo | Luke and Christianson, 2018 | Word |  | Short passages (2.5 sentences) | Visual | 2,659 | 84.00 | 0.93 |
| Beh | Eye-tracking | MECO-En | Siegelman et al., 2022 | Word |  | Passages (8.25 sentences) | Visual | 2,096 | 40.13 | 0.87 |
| Beh | Eye-tracking | ZuCo-2 | Hollenstein et al., 2020 | Word |  | Sentences | Visual | 6,461 | 17.44 | 0.76 |
| Beh | Eye-tracking | UCL<sub>ET</sub> | Frank et al., 2013 | Word |  | Sentences | Visual | 1,726 | 42.00 | 0.84 |
| Beh | SPR | UCL<sub>SPR</sub> | Frank et al., 2013 | Word |  | Sentences | Visual | 1,316 | 71.11 | 0.22 |
| Beh | SPR | Brown | Smith and Levy, 2013 | Word |  | Passages | Visual | 5,862 | 19.05 | 0.39 |
| Beh | SPR | NatStor<sub>SPR</sub> | Futrell et al., 2021 | Word | ✓ | Stories (48.5 sentences) | Visual | 8,779 | 59.63 | 0.70 |
| Beh | Maze | NatStor<sub>Maze</sub> | Boyce et al., 2023 | Word | ✓ | Stories (48.5 sentences) | Visual | 9,779 | 5.86 | 0.47 |
| Brain | N400 | UCL<sub>N400</sub> | Frank et al., 2015 | Word |  | Sentences | Visual (length-dependent presentation rate) | 1,505 | 24.00 | 0.22 |
| Brain | BOLD | Wehbe2014 | Wehbe et al., 2014 | TR |  | Chapter | Visual (fixed presentation rate) | 1,211 | 8.00 | 0.16 |
| Brain | BOLD | NatStor<sub>fMRI</sub> | Shain et al., 2022a | TR | ✓ | Stories (48.5 sentences) | Auditory | 1,516 | 27.18 | 0.61 |
| Brain | BOLD | Pereira2018 | Pereira et al., 2018 | Sentence | ✓ | Passages (4 sentences) | Visual | 627 | 7.84 | 0.28 |
| Brain | BOLD | Tuckute2024 | Tuckute et al., 2024b | Sentence |  | Sentences | Visual | 2,000 | 6.25 | 0.22 |

In terms of the LMs, we considered the original GPT model and four GPT-2 model variants (GPT-2_124M_, GPT-2_355M_, GPT-2_774M_, GPT-2_1.5B_). Additional details on the models and the datasets can be found in the Methods (§5.2, Table 1).

### 2.1 Behavioral responses reflect bottlenecking of representations into low-dimensional predictors

The results are summarized in Figure 3. Across all behavioral datasets, responses are primarily driven by surprisal, frequency, and length. Embedding-based regressors capture little additional variance, suggesting that behavioral responses are mostly determined by cognitive effort.

**Figure 3:**
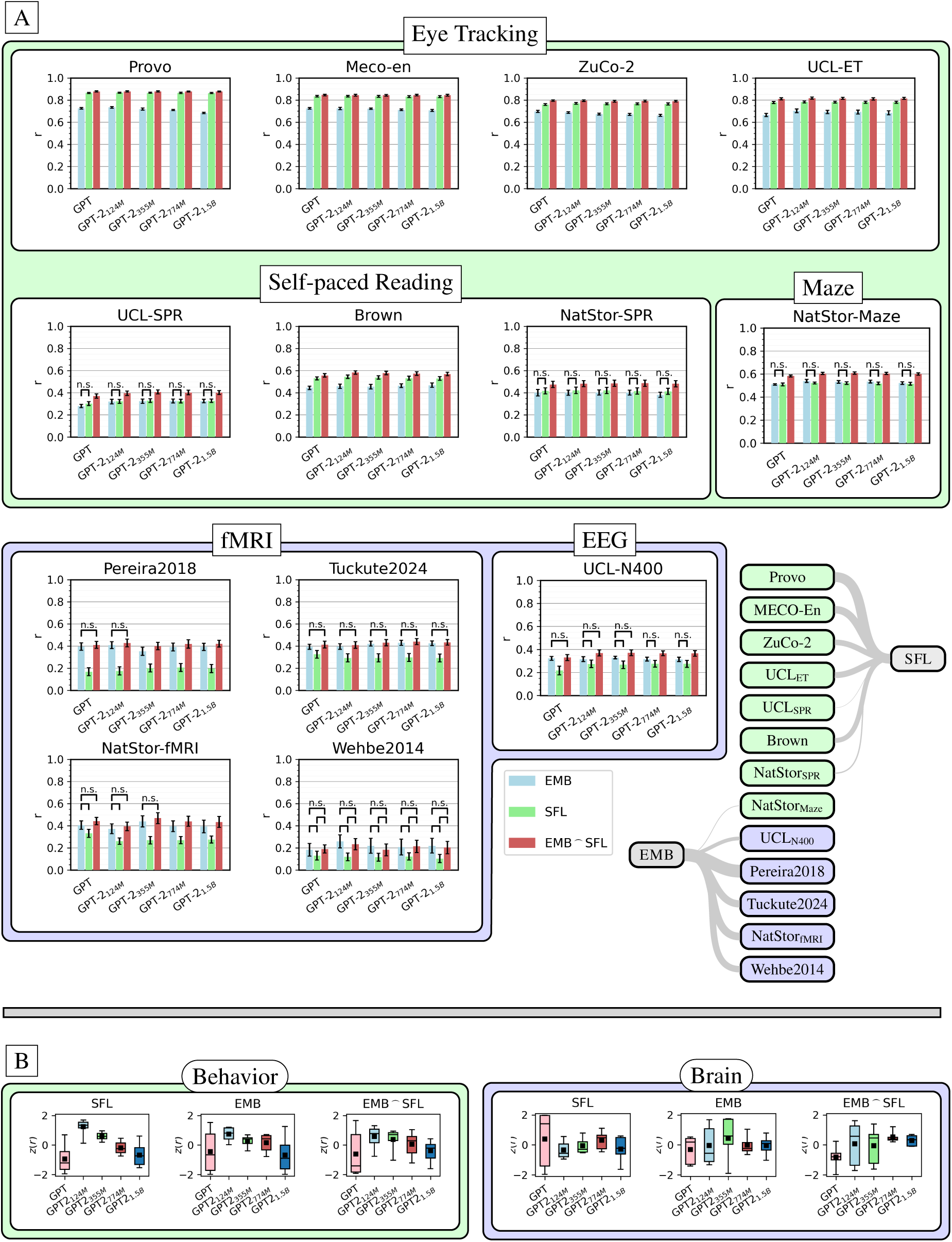
**A**: Graphical summary of the results obtained across datasets and models. The height of the bars indicates the average *r* obtained in the 10 folds, and the error bars report the standard error of the mean across folds. Given that the majority of the comparisons (EMB, SFL, EMB_⌢_SFL) were statistically significant, for readability we annotate the non-significant comparisons. The diagram on the right summarizes the comparison between EMB and SFL. The gray lines connect the behavioral datasets (green oblongs) and neural datasets (blue oblongs) to their best predictor among SFL and EMB; the width of the line is proportional to the difference between the results obtained with SFL and EMB. The diagram shows that SFL is generally associated with higher levels of predictivity for the behavioral datasets, whereas EMB achieves better results for the neural datasets. **B**: Model-focused summary of the results, aggregated across all behavioral datasets (left) and all neural datasets (right). The *r* values obtained with the different models in each dataset are *z*-scored. The black squares specify the mean value of the results obtained by each model and the horizontal lines indicate the median.

For the eye-tracking data using our primary measure (gaze duration, see Appendix A.4 for a replication with first fixation duration and total fixation duration), the general predictivity of SFL is higher than the one achieved by EMB (see Appendix A.3 for the individual contributions of surprisal, frequency, and length, as well as all pairwise combinations for all datasets). The average difference in predictivity between SFL and EMB across datasets and model sizes is 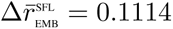. The concatenation of EMB to SFL (SFL^⌢^EMB) improves the results by a statistically significant but numerically small amount 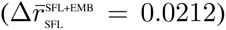. From a qualitative perspective, the results obtained across the four eye-tracking datasets are extremely consistent, with higher performance obtained by SFL^⌢^EMB, followed by SFL and lastly EMB alone. The results obtained for the self-paced reading datasets are qualitatively aligned with this trend, with SFL outperforming EMB (although by a smaller amount; 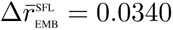, and significantly so only in Brown). In this case, however, EMB and SFL appear to provide more complementary information, as the addition of EMB to SFL increases the model fit by a relatively larger margin 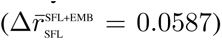. NatStor_Maze_ is the only behavioral dataset where EMB numerically outperforms SFL, although the difference is not statistically significant for any of the five models 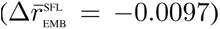; similar to what we found for NatStor_SPR_, SFL^⌢^EMB outperforms SFL 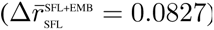.

Across the behavioral datasets, the best results are, on average, obtained by GPT-2_124M_—the smallest GPT-2 variant we consider—both when employing SFL and EMB representations, and more generally, we find that smaller models better account for the behavioral correlates of language comprehension. This “inverse scaling” trend replicates the previous finding that smaller models better account for human reading times (Oh and Schuler, 2022, 2023; de Varda and Marelli, 2023; Shain et al., 2024; Klein et al., 2024; Figure 3, panel A, bottom left).

### 2.2 Brain responses reflect rich linguistic content beyond processing effort

The results obtained with brain data are radically different: across all brain datasets, embedding-based predictors capture far more variance in brain activity than surprisal, frequency, and length.

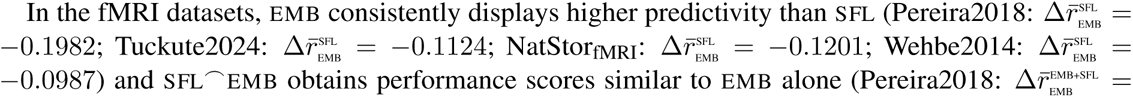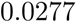. Similar results are obtained with the ERP dataset, where EMB numerically outperforms SFL in predicting N400 amplitudes 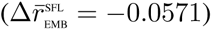. We wish to stress the consistency of the pattern across measures and datasets, although the pairwise comparisons between EMB and SFL do not generally reach statistical significance in UCL_N400_ and in Wehbe2014, due to the high variability in the *r* values obtained across folds. Our qualitative results hold even when using high-dimensional non-linear projections of SFL that match EMB in dimensionality, indicating that the advantage of EMB in predicting brain responses is not an artifact of its dimensionality (see Appendix A.6). Note also that the asymmetry between the behavioral and the neural results cannot be simply attributed to a difference in the experimental stimuli, as a subset of our datasets (the UCL corpus and NatStories) comprises both behavioral and brain responses to the same set of sentences; the UCL corpus further includes responses at the same temporal resolution (word-by-word). This characteristic of the experimental setup strengthens the conclusion that the difference is rooted in the nature of the dependent measures themselves.

Concerning model comparison, the trend moves in the opposite direction from the behavioral results: within the GPT-2 family, larger models tend to outperform their smaller counterparts, especially with the SFL predictor (see Figure 3, panel B, bottom right). Furthermore, with SFL and SFL^⌢^EMB, the smallest GPT-2 model (GPT-2_124M_) obtains the lowest performance within the GPT-2 family in predicting brain responses. This pattern aligns with previous findings that larger-capacity neural architectures obtain better brain encoding performance (Schrimpf et al., 2021; Antonello et al., 2023), highlighting the representational richness of linguistic representations in the human brain.

The diagram in Figure 3 (panel A, bottom right) graphically summarizes the comparison between the predictive power of SFL and EMB across the considered datasets; it clearly shows that behavioral measures are more strongly associated with the traditional predictors of cognitive effort, whereas neural responses are better predicted by model embeddings.

The differences between behavioral and brain responses to language further emerge when comparing the individual predictivity of the three components of SFL: behavioral responses—and especially eye movements during reading—are dominated by the context-independent properties of frequency and length, with surprisal playing a secondary role (Appendix A.3). Conversely, surprisal is the strongest predictor of brain responses, followed by frequency and length.

## 3 Discussion

In this paper, we showed that a low-dimensional, interpretable predictor quantifying cognitive effort—a combination of surprisal, word frequency, and length (SFL)—excels at capturing variance in behavioral measures of incremental language processing. This predictor outperforms information-rich embedding representations in explaining eye-tracking and self-paced reading data, and adding EMB to SFL yields only minor gains. This pattern suggests that the behavioral effects of model-derived representations are largely mediated by a small set of cognitively motivated features related to processing difficulty.

The stronger predictivity of SFL for behavioral data is particularly striking given that its dimensionality is more than two orders of magnitude lower than EMB. This disproportionate predictive power relative to dimensionality is in line with the long-standing interest that these predictors have received in psycholinguistic research over the last few decades, especially in studies that use eye-tracking during reading (see for instance Just and Carpenter, 1980; Rayner et al., 1996; Rayner and Duffy, 1986; see Rayner, 1998 for a review). Indeed, the importance of these features in explaining reading behavior has led researchers to label them as the “The Big Three” of lexical processing (Clifton Jr et al., 2016). Their influence on processing cost is well motivated from the perspective of a rational information processor. The length of a word is proportional to the number of letters or sounds that need to be recognized; thus, longer words are expected to more strongly tax the early stages of word processing. Word frequency approximates the number of encounters with a certain word form; frequency effects can thus be interpreted as the outcome of a word learning process and reflecting lexical processing (Brysbaert et al., 2018; Berzak and Levy, 2023) or as reflecting general sensitivity of the human parser to local statistics (Goodkind and Bicknell, 2021). Then, word predictability effects captured by surprisal are predicted by the view of the language processor as an optimal incremental comprehension system that makes efficient use of the linguistic context (Levy, 2008; Fernandez Monsalve et al., 2012; Smith and Levy, 2013; Kuperberg and Jaeger, 2016; Berzak and Levy, 2023; Shain et al., 2024). Our findings broadly support the bottlenecking hypothesis (Levy, 2008): although EMB provides a minor improvement in some cases, the size of this improvement is generally small. These results are therefore best understood as evidence that behavioral responses reflect a highly compressed encoding of linguistic information, sensitive primarily to the most salient dimensions of processing difficulty.

For the neural responses during language processing, high-dimensional embedding representations produced by language models are the best predictors of both fMRI responses and the N400 ERP component, consistently outperforming SFL across datasets. Although effort-related components predict neural responses to a non-trivial extent, a substantial portion of additional variance is accounted for by embeddings. In the context of prior work, this pattern reconciles two observations: robust neural sensitivity to processing difficulty (e.g., Federmeier and Kutas, 1999; Frank et al., 2015; DeLong et al., 2005; Federmeier et al., 2007; Henderson et al., 2016; Shain et al., 2020; Heilbron et al., 2022) and to higher-level aspects of linguistic representation, including the construction of linguistic meaning (e.g., Huth et al., 2016; Deniz et al., 2019; Tuckute et al., 2024b, 2025). Our results suggest that effort-related effects on brain responses reflect a constrained readout of richer contextual representations, capturing only a subset of the information encoded in the neural signal. Consistent with this view, adding SFL to EMB provides little improvement, indicating that embeddings subsume the predictive contribution of the effort-related predictors in neural responses.

Two further observations support the interpretation that neural responses are sensitive to richer contextual representations. First, within the GPT-2 family, larger models yield better fits to brain data (see also Schrimpf et al., 2021; Antonello et al., 2023), in contrast to the inverse scaling trend observed for behavior. Second, when decomposing SFL, surprisal emerges as the dominant predictor of brain responses, and frequency and length play a secondary role—again, the opposite of the pattern found in behavioral data, where context-independent properties show the strongest influence. Surprisal is the only context-dependent component of SFL, so the contrast between brain and behavioral data reappears within SFL: brain responses are most strongly modulated by the context-sensitive predictor, whereas behavioral responses mostly reflect the context-invariant ones. Together, these results suggest that, perhaps unsurprisingly, neural measures provide access to the rich, context-dependent representations of meaning that underlie language comprehension.

Why, then, do behavioral measures show limited sensitivity to rich, high-dimensional linguistic representations, despite clear evidence that such information is encoded in the brain? The reason, we argue, is that the timescale of reading differs from that of deep semantic and syntactic integration. During “default” fluent reading of ordinary texts, which typically do not contain anomalies or exceptionally complex structures, the eyes move quickly and often leave a word before it has been fully processed (Clifton Jr et al., 2016). The mechanisms that support deep understanding of the content are too slow to guide when and where the eyes move. Overt reading behavior is therefore driven mainly by shallow, rapidly available signals (SFL) compared to fully formed linguistic representations (EMB). This view is instantiated in the E-Z Reader framework (Reichle et al., 1998, 1999, 2003), which posits that word identification unfolds in two stages: an early stage that provides a fast estimate of whether word identification is imminent and initiates saccade programming, and a later stage that supports fuller linguistic processing but does not directly control overt eye movements. Under this account, higher-order syntactic and semantic representations can influence comprehension without being reliably expressed in moment-to-moment reading behavior. Broadly, this perspective suggests that behavioral measures obtained during fluent reading constitute a bottlenecked readout of language processing, selectively reflecting those aspects of linguistic computation that are available early enough to influence overt behavior. The E-Z Reader account was developed for eye movements, but a similar logic may apply to self-paced reading and the Maze paradigm, where the motor response is also produced quickly and may precede full linguistic integration.

It is important to note that linguistic effects that cannot be reduced to shallow factors such as SFL are well documented, particularly in the domain of syntax and in cases where comprehension is disrupted (e.g., garden-path sentences, Frazier and Rayner, 1982; Ferreira and Henderson, 1991; long-distance syntactic dependencies, Grodner and Gibson, 2005; Warren and Gibson, 2002; Boston et al., 2008; Bartek et al., 2011). Under such conditions of unusually high processing complexity, reading times may reflect not only fast, early estimates of lexical processing difficulty (which are well approximated by SFL), but also other costs related to memory-intensive linguistic operations or re-analysis of preceding input (some of which might be captured by EMB). And indeed, the behavioral bottleneck discussed here should be understood as a property of “default” fluent reading, whereas higher-order processes may intervene in eye-movement control only when there is a need for re-analysis (see also Reichle et al., 1998). Support for this interpretation comes from the study of memory effects in online language processing behavior, which can be robustly detected with experimentally constructed materials that manipulate the length of syntactic dependencies (Grodner and Gibson, 2005; Boston et al., 2008; Bartek et al., 2011), but do not emerge when participants read typical texts (Demberg and Keller, 2008; Shain et al., 2016, see Gibson, 2025 for discussion).

This line of reasoning also helps explain one pattern in our own behavioral results. The contrast between effort-and embedding-based predictors is less pronounced in the two behavioral NatStor datasets (Maze and self-paced reading), which contain many unusual syntactic constructions (Futrell et al., 2018) and thus deviate from reading behavior that characterizes more typical texts. In these materials, behavioral responses likely reflect not only shallow lexical processing difficulty, but also additional costs associated with complex syntactic integration and re-analysis. Some of these additional costs are captured by surprisal, which contributes more to behavioral predictivity in NatStor than in other datasets (Appendix A.3). However, surprisal has been shown to systematically fail to account for the costs of syntactic integration and re-analysis (Arehalli et al., 2022; Van Schijndel and Linzen, 2021; Hahn et al., 2022; Wilcox et al., 2021; Huang et al., 2024). In those cases, embedding representations may capture the linguistic components that underlie these costs, predicting additional variance in language processing behavior over SFL. Future work should extend our approach to experimentally controlled, non-naturalistic materials to understand how and when richer representations affect both behavioral and neural responses.

## 4 Conclusion

In this study, we compared EMB and SFL in terms of how well they predict behavioral and brain measures of language processing. We showed that the relative fit of these two predictors critically depends on the outcome variable: behavioral responses are best accounted for by SFL, whereas brain responses are better captured by EMB. Language processing behavior thus reflects a compressed bottleneck over a few theoretically motivated properties of the input, whereas neural data retain access to a richer, high-dimensional representational space.

The analyses in this paper focused primarily on predictive performance. Beyond predictivity, EMB and SFL also differ in the interpretative commitments they invoke. The functional significance of surprisal, frequency, and length can be explained in terms of rational information processing, and, in the case of surprisal, in algorithmic proposals specifying why the human processor should be sensitive to word predictability, for instance in terms of Bayesian inference (Marr, 1982; Bicknell and Levy, 2010). By contrast, convergence between model and human representations in the EMB framework is often given a teleological reading (Cichy and Kaiser, 2019) and thus situated at the computational level of description (Marr, 1982): brains and models would align because they are both optimized for prediction (Schrimpf et al., 2021; Caucheteux and King, 2022; cf. Antonello and Huth, 2024) or deeper language understanding (Aw and Toneva, 2022).

A second axis of comparison beyond predictivity concerns model complexity and parsimony. Our analyses deliberately set aside one consideration that might otherwise guide model selection: the complexity of the predictive model itself. In philosophy of science, when competing models yield comparable predictions, parsimony is often invoked as a guiding principle (Ockham, 1495; cf. Dubova et al., 2025); furthermore, common approaches in model comparison involve a penalty for complexity (Akaike, 1998). In this respect, the comparison between EMB and SFL reflects a trade-off that characterizes many modeling efforts in science: the balance between predictivity on one hand and simplicity and interpretability on the other (Tuckute et al., 2024b). As models of behavioral data, SFL representations satisfy both desiderata: they are not only highly predictive of reading behavior, but also low-dimensional and interpretable. As such, they offer a compelling explanatory framework for the behavioral correlates of comprehension difficulty. When modeling brain data, however, the situation is more complex. EMB obtains strong predictivity, but at the cost of increased model complexity; SFL is interpretable, but does not match the predictive capacity of EMB.

Nonetheless, the high complexity of EMB should not make us think of it as a less valuable tool for understanding how language is encoded in the brain. Rather, it suggests that understanding brain responses to language may require engaging with the full representational richness encoded in language model embeddings. Future work may leverage post-hoc analytical tools to decompose the embedding space and isolate structured, interpretable components that drive its alignment with neural responses (Tuckute et al., 2024b).

## 5 Methods

### 5.1 Models

In our analyses, we considered five GPT variants, namely the original GPT model (Radford et al., 2018) and four models in the GPT-2 family with parameter size spanning from 124M to 1.5B (GPT-2_124M_, GPT-2_355M_, GPT-2_774M_, GPT-2_1.5B_; Radford et al., 2019); we employed the same models to derive surprisal estimates and embedding representations. In the case of EMB, word-level representations were obtained by averaging over sub-word tokens. Surprisal values were instead summed over sub-word tokens. In our analyses, we generally considered the embeddings and surprisal values relative to *w*_i_; those corresponding to previous words were included in the analyses of neural and behavioral variables that are well-known to be responsive to spill-over effects, namely fMRI responses and self-paced reading times.

### 5.2 Datasets

Our analyses were performed on four eye-tracking, three self-paced reading, one Maze, one ERP, and four fMRI datasets. We provide a summary of the datasets we considered in Table 1. Given the high number of resources we considered in our analyses, we briefly describe each dataset individually in the Supplementary Information (A.1); we further redirect the reader to the original publications for additional details.

### 5.3 Pre-processing

#### 5.3.1 Behavioral data

We analyzed the behavioral datasets following a consistent pre-processing pipeline. For eye-tracking data, we considered first-pass gaze duration times; words that were not fixated were counted as having a gaze duration time equal to zero. (Similar results were obtained using first fixation duration and total fixation duration; Appendix A.4). Sentence-initial words were excluded from the analyses; apart from this criterion, we did not apply any other filtering unless stated otherwise. Word-level gaze duration times, self-paced reading reaction times, and Maze reading times were obtained by averaging responses across participants. Given that self-paced reading times are well-known to be sensitive to spill-over effects, the properties of the words *w*_i*−*2…i_ were retained for the analyses. We did not include spill-over effects for eye-tracking data and Maze reading times since those two measurements have high temporal resolution and produce small spill-over effects, if any (Forster et al., 2009; Boyce et al., 2023; Wilcox et al., 2023).

#### 5.3.2 Neural data

##### fMRI datasets

The fMRI datasets were analyzed differently depending on whether responses were recorded for continuous stories (Wehbe2014, NatStor_fMRI_) or individual sentences (Pereira2018, Tuckute2024). For the stories, functional data were acquired with repetition time (TR) equal to 2 seconds, and words were either visually presented at a fixed interval of 0.5 seconds (Wehbe2014) or recorded by a speaker, thus pronounced at a variable interval. Thus, to obtain TR-level predictors, both EMB and SFL were downsampled by averaging over four words (Wehbe2014) or the number of words pronounced during the TR interval (NatStor_fMRI_). In the case of NatStor_fMRI_, word-by-word timestamps necessary to temporally align EMB and SFL with the fMRI responses were extracted with the speech recognition model Whisper-timestamped (Radford et al., 2023; Louradour, 2023). Following Aw and Toneva (2022), we employed as predictors the concatenation of the SFL and EMB values relative to the four previous TRs, to account for the lag in the hemodynamic response recorded with fMRI. For the fMRI datasets including responses to sentences, sentence-aggregated SFL and EMB regressors were obtained by averaging over word-level predictors. Across all four fMRI datasets, we considered as our core response variable the average activity across the language network (see also Tuckute et al., 2024b; de Varda et al., 2025). Aggregated responses were obtained by first averaging across the voxels within (functional) regions of interest (fROIs), then across fROIs, and lastly across participants, to obtain a univariate response, necessary for meaningful comparisons with the intrinsically univariate behavioral responses. In three datasets (NatStor_fMRI_, Tuckute2024, Pereira2018), regions of interest were identified functionally through a localizer task, whereas in the Wehbe2014 dataset, regions of interest were identified anatomically (see §A.2.2).

##### ERP dataset

We considered N400 responses from the UCL_N400_ corpus, where words were visually presented at a word-length dependent presentation rate, considering N400 amplitudes from 12 centroparietal electrodes. EEG responses were filtered following the exclusion criteria described in Frank et al. (2015); our analyses were based on the pre-computed word-by-word N400 amplitudes. Word-level N400 responses were obtained by averaging across participants. Words adjacent to punctuation signs were excluded after noticing during preliminary analyses that punctuation had a prominent effect on N400 responses, possibly due to wrap-up effects. Note however that the direction of the results reported in the paper was consistent regardless of the exclusion of punctuated words.

### 5.4 Analyses

With the EMB approach, an important concern is to avoid overfitting. Indeed, the number of parameters of the regression models is typically larger than the number of observations. This complication must be contrasted with cross-validation and regularization. We used ridge regularization, due to its computational efficiency (Tikhonov, 1963), and 10-fold cross-validation throughout our analyses, with the exception of NatStor_fMRI_, where we held out one story at a time (9 folds), so that no story contributed to both the training and the test set. We selected the penalty term *α* in each training fold through nested 5-fold cross-validation, considering 13 candidate penalties ranging from 10*^−^*^6^ to 10^6^ (logarithmically spaced). Predictors and responses were standardized within each training fold (applying the training-fold statistics to the test fold), so that the ridge penalty applies uniformly across features regardless of their scale. When fitting the concatenation SFL^⌢^EMB, we employed banded ridge regression (Nunez-Elizalde et al., 2019; La Tour et al., 2022) which assigns a separate penalty to each feature block, necessary because the features are on different scales. We adopted a sequential train-test split since with shuffled train-test splits it is possible to achieve high predictivity by simply relying on the temporal autocorrelation of the signal (Kauf et al., 2024; Hadidi et al., 2026). In the Pereira2018 dataset, sentences are grouped into topically coherent passages; we therefore held out whole passages at each fold, so that sentences from the same passage never appeared in both the training and the test set. As a performance metric, we employed the Pearson correlation *r* between the predictions of the regression model and the actual values of the behavioral or neural measure of interest, averaging across folds. To assess statistical significance, we used non-parametric paired-samples permutation tests. Prediction scores from each cross-validation fold were permuted between the two alternatives (e.g., SFL and EMB), and the test statistic (the difference in mean *r* values) was recomputed for all 2^n^ possible permutations. The *p*-value was defined as the proportion of permuted differences equal to or greater than the observed one.

### 5.5 AI disclosure

Claude Opus 4.8–5 was used to assist with writing and debugging the analysis code, and for light copy-editing. All code was tested and validated by the authors, who take full responsibility for the analyses and results reported here.

### 5.6 Data and code availability

All the code and data employed in this study are publicly available.^1^

## Acknowledgements

We thank Guangyuan Jiang for suggesting and exploring the control analysis discussed in Appendix A.6, James Michaelov for useful suggestions on the framing, and all members of the Computational Psycholinguistic Lab for feedback and discussion. AGdV was supported by the K. Lisa Yang ICoN Center Fellowship. EF was supported by research funds from the McGovern Institute for Brain Research, the Simons Center for the Social Brain, and the MIT Siegel Family Quest for Intelligence. YB and RL were supported by BSF grant 2024151.

## **A** Supplementary information

### **A.1** Data description

#### A.1.1 Provo

The Provo corpus (Luke and Christianson, 2018) comprises eye-tracking data from 84 native speakers of American English. Participants read 55 short passages from various sources, including news articles, science magazines, and public-domain fiction.

#### **A.1.2** MECO-En

The Meco corpus (Siegelman et al., 2022) is a collection of eye-movement data in 13 languages. For the purpose of this study, we focused on the English component of the corpus (MECO-En). Participants engaged in a naturalistic reading task, and were presented with 12 texts consisting of encyclopedic entries on a handful of topics. MECO-En includes data from 46 participants.

#### **A.1.3** ZuCo-2

The ZuCo-2 corpus (Hollenstein et al., 2020) is a dataset of simultaneous eye-tracking and electroencephalography recordings including data from 18 participants. In this study, we focused on the eye-tracking records. The resource comprises chronometric data from 349 sentences presented in a normal reading paradigm, and 390 presented in a task-specific paradigm, where the participants were instructed to actively search for certain semantic relations. As our main interest is natural reading, we conducted our analyses on the former portion of the dataset.

#### A.1.4 UCL_ET_

The UCL_ET_ corpus (Frank et al., 2013) is a resource consisting of eye movement data on 205 sentences read by 43 participants. The sentences were extracted from publicly available unpublished novels, selecting those that contained only high-frequency words and could be interpreted out of context.

#### A.1.5 UCL_SPR_

The self-paced reading component of the UCL reading corpus (Frank et al., 2013) includes chronometric data from 117 participants reading 361 sentences. To increase the comparability with the eye-tracking component of the corpus, we only considered the 205 sentences shared with UCL_ET_. Words were presented at the center of the screen, and the current word was replaced by the next one at each subsequent keypress.

#### **A.1.6** Brown

The Brown corpus (Smith and Levy, 2013) is a resource of moving-window self-paced reading times collected from 35 participants, who read a varying portion of the corpus. The passages were drawn from the Brown corpus of American English.

#### **_A.1.7_** NatStor_SPR_

The NatStor_SPR_ corpus (Futrell et al., 2021) includes moving-window self-paced reading times collected from 181 participants. The corpus consists of 10 stories developed by modifying publicly available texts to contain several rare syntactic structures (e.g., subject-and object-extracted relative clauses, clefts, topicalized structures). The syntactic editing of the texts aimed to maintain their meaning, and conserve a high degree of comprehensibility. Most participants read 5 out of the 10 stories.

#### **_A.1.8_** NatStor_Maze_

The NatStor_Maze_ corpus (Boyce et al., 2023) is a resource of Maze reading times collected on the same texts as NatStor_SPR_. In the Maze task, reaction times are recorded while participants iteratively choose between two words, presented simultaneously: a correct word that continues the sentence, and a distractor string which does not. In this dataset, an “error-correction” variant of the Maze task was employed, in which when participants make an error, they see an error message and are asked to select again the correct word. Distractors were automatically generated. NatStor_Maze_ includes Maze reading time data from 63 participants; each participant read one of the 10 stories. Maze data were cleaned following the exclusion criteria described by Boyce et al. (2023).

#### **_A.1.9_** UCL_ERP_

The UCL_ERP_ corpus (Frank et al., 2015) includes amplitudes of six ERP components (N400, P600, and ELAN, LAN, PNP, and EPNP—four other ERP components which have received comparatively little attention in the literature). In this work, we focused on N400 amplitudes, which have been shown to be strongly responsive to surprisal in this dataset (Frank et al., 2015; de Varda et al., 2024). The EEG recordings were collected from 24 participants reading 205 sentences. Words were serially presented at the center of the screen at a word-length dependent presentation rate. N400 amplitudes were operationalized as the average voltage of 12 centro-parietal electrodes 300-500 ms after the presentation of each word (electrodes sites 1, 14, 24, 25, 26, 29, 30, 31, 41, 42, 44, 45).

#### **A.1.10** Wehbe2014

The Wehbe2014 dataset (Wehbe et al., 2014) comprises fMRI recordings of 8 participants reading the ninth chapter of the book *Harry Potter and the Sorcerer’s Stone* (Rowling, 2000). The chapter’s words (N = 5,176) were displayed sequentially at a fixed presentation rate of 0.5 seconds; they were split into four runs of comparable length. Functional imaging data were sampled with TR of 2 seconds.

#### **_A.1.11_** NatStor_fMRI_

The NatStor dataset contains fMRI recordings from participants who listened to naturalistic stories (approximately 5 minutes each). Data collection took place in the Fedorenko lab between 2014 and 2020. Initial results were reported in Blank et al. (2014), and subsequent analyses based on the expanded dataset appeared in several studies (Paunov et al., 2022; Shain et al., 2020, 2022b; Sueoka et al., 2024; Wehbe et al., 2021). The dataset in its current form has been used by de Varda et al. (2025). The dataset includes 9 stories: 8 were edited to incorporate low-frequency syntactic constructions while preserving naturalness for native speakers (see Futrell et al., 2021), and 1 (an expository piece titled “Tree”) was adapted from Wikipedia (Paunov et al., 2022). The number of participants per story ranged from 7 to 70, with a mean of 27.18.

#### **A.1.12** Pereira2018

The Pereira2018 dataset (Pereira et al., 2018; Experiments 2 and 3) includes fMRI recordings from 10 unique participants (9 in Experiment 2 and 6 in Experiment 3, with 5 participating in both) reading 627 sentences (384 in Exp. 2, 243 in Exp. 3), grouped into passages of three to four sentences. All passages were encyclopedic texts providing basic information about a concept. Each participant saw each sentence between 1 and 4 times, with the majority seeing each sentence 3 times. The two sets of materials were constructed independently, and each spanned a broad range of content areas. Sentences were 7–18 words long in Experiment 2, and 5–20 words long in Experiment 3. The sentences were presented visually on a screen one at a time for 4 seconds (followed by 4 seconds of fixation, with additional 4 seconds of fixation at the end of each passage).

#### **A.1.13** Tuckute2024

The Tuckute2024 dataset (Tuckute et al., 2024b) comprises fMRI data from 10 participants who read a total of 2,000 sentences (with 6.25 responses per sentence, on average). These sentences, all six words in length, were sourced from written and transcribed spoken corpora to cover a wide range of content and styles. Sentences were displayed visually on a screen for 2 seconds each, followed by a 4-second fixation period. Data were collected over two or three separate scanning sessions.

#### **A.2** fMRI pre-processing and first-level analysis

The Pereira2018, Tuckute2024, NatStor_fMRI_ datasets were pre-processed following the same procedure (§A.2.1); across all three datasets, responses were recorded from five functional regions of interest (fROIs) defined in individual participants through a “localizer task” (§A.2.2). We then averaged the responses across the voxels within each of the five fROIs, then across fROIs, and lastly, across participants, to obtain a univariate response for each dataset (thus comparable with the behavioral responses). The Wehbe2014 dataset did not include fROI information since no localizer was used during data collection.

#### **A.2.1** Pre-processing (shared between Pereira2018, Tuckute2024, NatStor_fMRI_)

Data preprocessing was performed with SPM12 (using default parameters, unless specified otherwise) and custom scripts in MATLAB. Preprocessing of anatomical data included normalization into a common space (Montreal Neurological Institute [MNI] template) and segmentation into probabilistic maps of the gray matter (GM), white matter (WM), and cerebro-spinal fluid (CSF). A GM mask was generated from the GM probability map and resampled to 2 mm isotropic voxels to mask the functional data. Preprocessing of functional data included motion correction (realignment to the mean image using second-degree b-spline interpolation), normalization (estimated for the mean image using trilinear interpolation), resampling into 2 mm isotropic voxels, and smoothing with a 4 mm full-width, half-maximum (FWHM) Gaussian filter.

#### **A.2.2** fMRI data modeling and definition of functional regions of interest (fROIs)

The language localizer data and the Pereira2018 and NatStor_fMRI_ were all modeled using the same setup (the setup was mostly similar in the Tuckute2024 dataset, with a few minor differences described in §A.2.3). Effects were estimated using a General Linear Model (GLM) in which each experimental condition was modeled with a boxcar function convolved with the canonical hemodynamic response function (HRF) (fixation was modeled implicitly, such that all timepoints that did not correspond to one of the conditions were assumed to correspond to a fixation period). Temporal autocorrelations in the BOLD signal timeseries were accounted for by a combination of high-pass filtering with a 128 seconds cutoff, and whitening using an AR(0.2) model (first-order autoregressive model linearized around the coefficient a=0.2) to approximate the observed covariance of the functional data in the context of Restricted Maximum Likelihood estimation (ReML). In addition to experimental condition effects, the GLM design included first-order temporal derivatives for each condition (included to model variability in the HRF delays), as well as nuisance regressors to control for the effect of slow linear drifts, subject-motion parameters, and potential outlier scans on the BOLD signal. Language fROIs were defined in individual participants (Fedorenko et al., 2010; Saxe et al., 2006) by combining two sources of information: (1) the participant’s activation map for the language localizer contrast (Fedorenko et al., 2010) and (2) group-level constraints (‘parcels’) that delineated the expected gross locations of activations and were sufficiently large to encompass the variability in the locations of individual activations (all parcels are available for download from https://www.evlab.mit.edu/resources-all/download-parcels). The language localizer, described in detail in Fedorenko et al. (2010), consisted of reading English sentences and lists of unconnected nonwords in a standard blocked design, with stimuli presented one (non)word at a time. The parcels were derived from a probabilistic activation overlap map using watershed parcellation, as described by Fedorenko et al. (2010), for the sentences *>* non-words contrast in 220 independent participants and covered extensive portions of the lateral frontal, temporal, and parietal cortices. Five language fROIs were defined (as the 10% of voxels with the highest t-values for the sentences *>* nonwords contrast) in the dominant hemisphere: three on the lateral surface of the frontal cortex and two on the lateral surface of the temporal and parietal cortex.

#### **A.2.3** fMRI data modeling – Tuckute2024

Effects were estimated using a GLM in which each experimental condition (i.e., each sentence trial) was modeled using GLMsingle (Prince et al., 2022). Using this framework, a general linear model (GLM) was used to estimate the beta weights that represent the BOLD response amplitude evoked by each individual sentence trial (fixation was modeled implicitly, such that all time points that did not correspond to one of the conditions (sentences) were assumed to correspond to a fixation period). For each voxel, the HRF that provided the best fit to the data was identified (on the basis of the amount of variance explained). The data were modeled using a fixed number of noise regressors (five) and a fixed ridge regression fraction (0.05) (these parameters were determined empirically using a joint data modeling and data evaluation framework; see Tuckute et al., 2024b). By default, GLMsingle returns beta weights in units of percent signal change by dividing by the mean signal intensity observed at each voxel and multiplying by 100. To mitigate the effect of collecting data across multiple scanning sessions, the beta values were z-scored session-wise per voxel.

#### **A.2.4** Pre-processing and modeling (Wehbe2014)

Functional data were preprocessed using SPM8 and custom MATLAB scripts. For each participant, preprocessing included realignment, slice timing correction, and co-registration of functional images to the anatomical scan, which was segmented into gray matter, white matter, and cerebrospinal fluid (CSF). The functional data were normalized to MNI space and smoothed with a 6 mm full-width at half maximum (FWHM) Gaussian kernel. Using PyMVPA (Hanke et al., 2009), the functional data were masked with the segmented anatomical image to exclude non-cortical voxels, including CSF. Temporal preprocessing included high-pass filtering at 0.005 Hz to remove low-frequency drifts and large-scale block effects, with the cutoff selected based on visual inspection of voxel time courses.

ROIs were defined on the basis of the AAL atlas (Tzourio-Mazoyer et al., 2002). See Wehbe et al. (2014) for additional details. Among those, we focused on regions that were considered in previous studies using this dataset, namely Posterior Temporal, Anterior Temporal, Angular Gyrus, Inferior Frontal Gyrus, Middle Frontal Gyrus, Inferior Frontal Gyrus (orbital part), and Posterior Cingulate (Aw and Toneva, 2022; Merlin and Toneva, 2024).

#### **A.3** Surprisal, frequency, and length

We report in Figure 4 the predictivity associated with surprisal (S), frequency (F), and length (L) taken alone as predictors of reading behavior and language-dependent brain responses. We further report the results obtained with all the possible feature subsets (SF, FL, SL, SFL). The reported results were obtained with GPT-2_355M_, as the model obtained solid average predictivity across the behavioral and brain datasets. In the datasets where spill-over effects were taken into account, the features of *w*_i_*, w*_i*−*1_ *. . . w*_i*−*N_ were jointly selected in the various combinations to avoid combinatorial explosion. The plots show that word length is the main determinant of gaze duration times, with little additional predictivity added by frequency and surprisal. With self-paced reading times, the predictivity of the three features is more homogeneous, but length remains the strongest predictor. NatStor_Maze_ is the only behavioral dataset in which surprisal is the strongest of the three predictors. In the brain datasets, surprisal has an edge over frequency and length across all datasets but Wehbe2014, where the performance obtained by frequency alone is slightly better. Across all neural datasets, word length exerts a reduced predictivity on brain responses.

**Figure 4:**
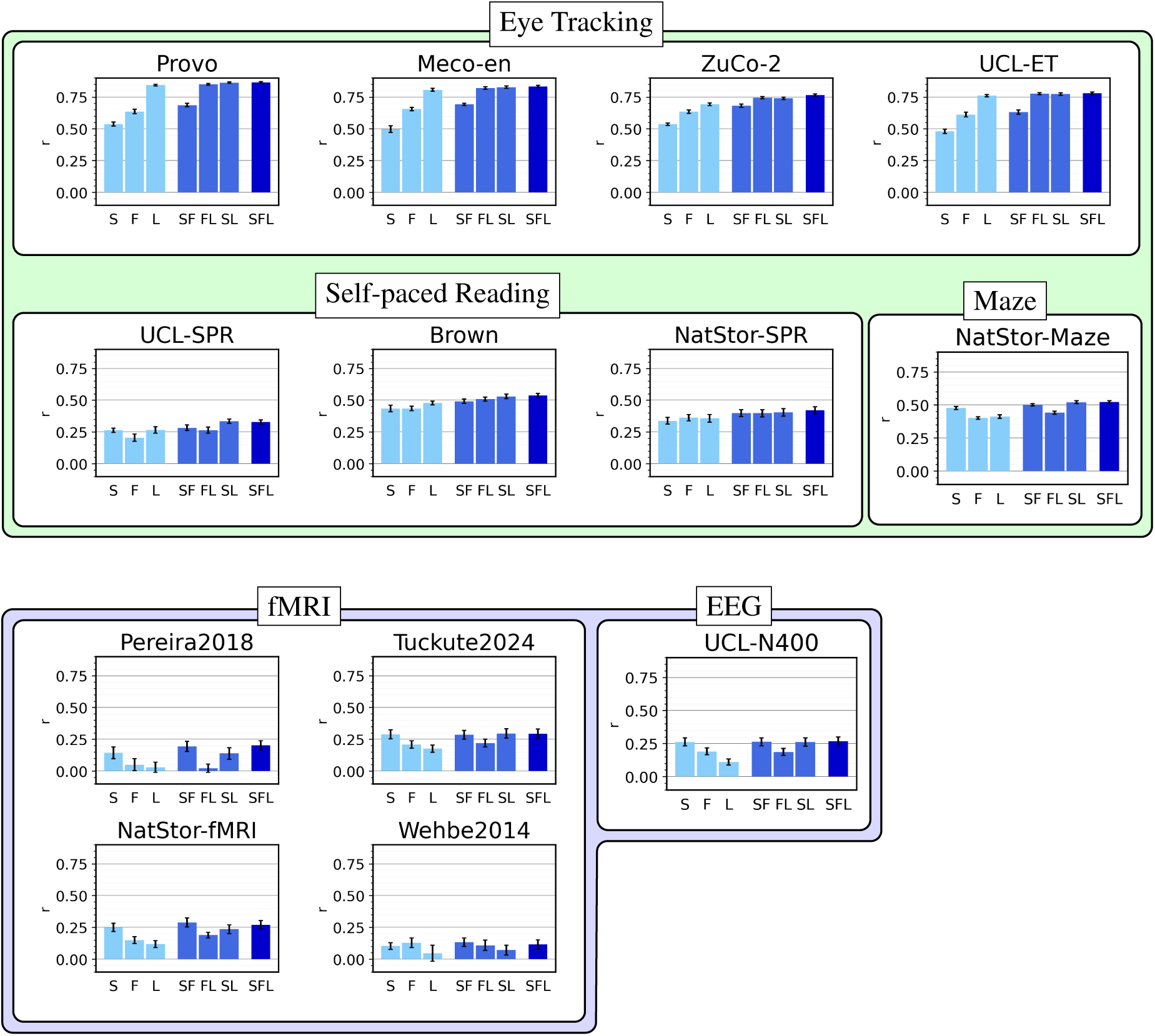
Predictivity of surprisal, frequency, and length with respect to the various neural and behavioral measurements; the models are fit with the predictors being analyzed individually (first three bars of each plot), with each two-predictor combination (second three bars), and with all three predictors (last bar). Length is associated with the highest predictivity in the eye-tracking datasets, whereas the three predictors obtain more homogeneous results in the self-paced and maze datasets. In the brain datasets, surprisal shows an advantage over the other predictors.

**Figure 5:**
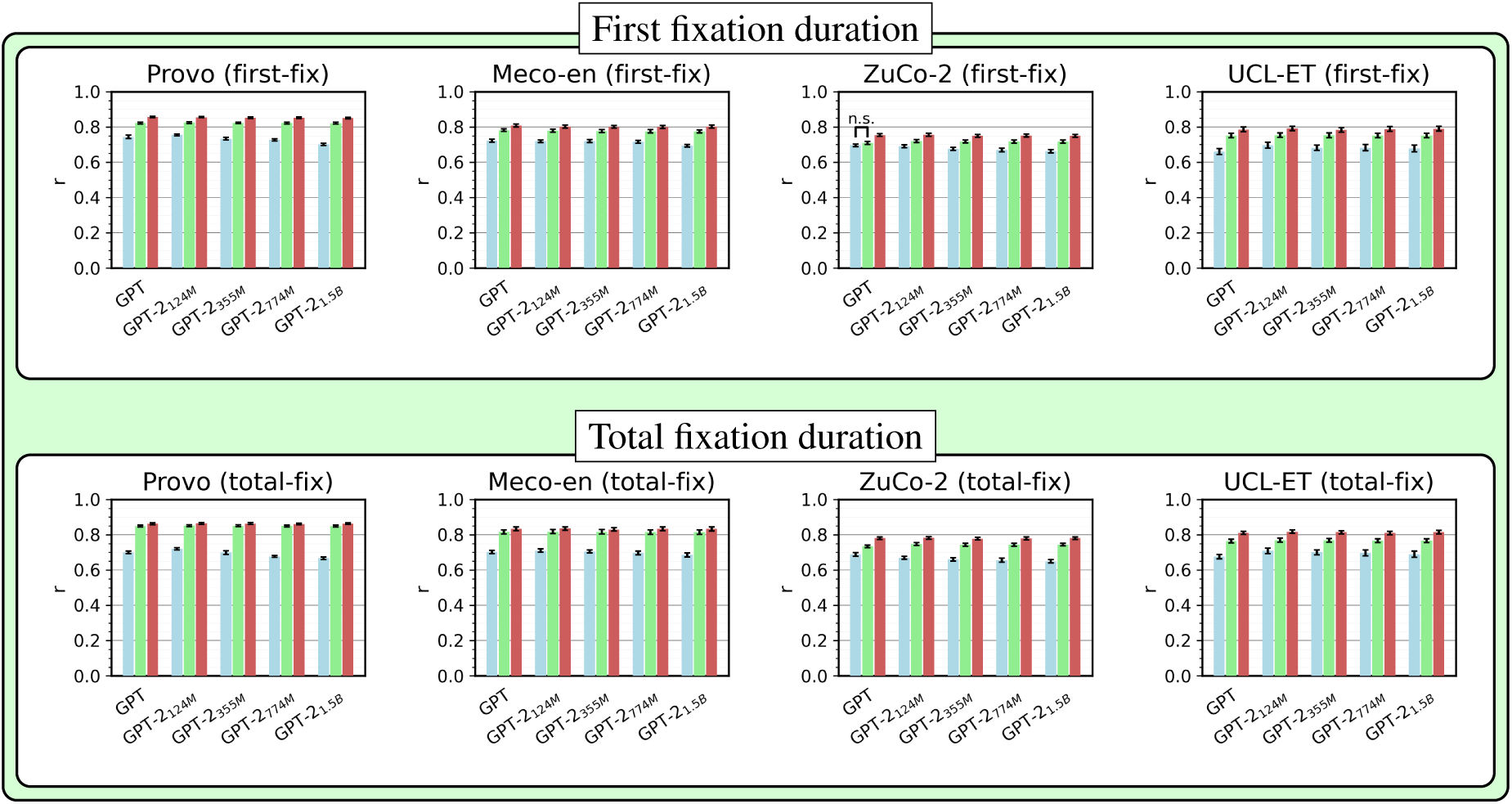
Predictivity of EMB, SFL, and EMB_⌢_SFL for first fixation duration and total fixation duration in the four eye-tracking datasets considered in the study. The general trends found in the main analyses considering gaze duration as the dependent variable (Figure 3, top) were largely replicated with the two additional fixation measurements.

#### **A.4** Additional fixation measures

In the main body of the paper, we presented the results obtained with gaze duration as the main dependent variable when analyzing eye-tracking data. Here, we replicated the analyses considering two additional measures, namely first fixation duration (the first single fixation on *w*_i_) and total fixation duration (the total amount of time the eyes spent on *w*_i_, including fixations returning to the word after having left it). Results confirmed our previous observation that on reading times data, SFL outperforms EMB in predicting word-by-word fixation times, and the concatenation of SFL^⌢^EMB yields reduced improvements over SFL alone.

#### **A.5** Layer-wise analyses

In the main analyses, we extracted embeddings from the final layer of each model. This choice maximizes comparability across datasets and is conservative in the case of brain data, where prior work has shown that intermediate layers often yield higher predictivity (Caucheteux and King, 2022; Antonello et al., 2023). For behavioral data, however, this choice may obscure the full predictivity of EMB: if embeddings from earlier layers better align with behavioral measures, then the gap in predictivity between EMB and SFL may be overestimated. Indeed, Tsipidi et al. (2026) found that representations from earlier layers better predict eye movement patterns, and Kuribayashi et al. (2025) documented that also surprisal values derived from earlier layers best predict reading times.

To evaluate this possibility, we re-extracted EMB representations from every layer of GPT-2_355M_, chosen because it is intermediate in size and obtains solid average predictivity on both brain and behavioral data (see also §A.3). Then, we compared each layer against SFL, which does not depend on the layer. Statistical significance was assessed with the same paired permutation test used throughout the paper (§5), but given the number of cells (13 datasets *×* 25 layers), we report differences that survive a Benjamini-Hochberg correction at *q* = .05 across all tests. The results are reported in Figure 7.

In the eye-tracking datasets, SFL remains the better predictor across layers: EMB never significantly outperforms it in Provo, MECO-En, or UCL_ET_, and does so only at layers 3–9 in ZuCo-2. In all four datasets EMB peaks very early (layer 4 in Provo, ZuCo-2, and UCL_ET_; layer 5 in MECO-En) and then declines monotonically. In the deepest layers of the network, SFL significantly outperforms EMB in every eye-tracking dataset (Provo, layers 7–24; MECO-En, 10–24; ZuCo-2, 14–24; UCL_ET_, 20–24). The representations that encode the most linguistic content are thus the ones that predict eye movements worst. This profile is consistent with the view that reading behavior is a shallow readout of language processing (§3). In the self-paced reading datasets the two predictors obtain similar predictivity in line with the main results (§2) apart from Brown, where SFL significantly outperforms EMB at layers 21–24. Like in the main results, NatStor_Maze_ is the only exception to the behavioral analysis, as EMB significantly outperforms SFL in most layers.

Finally, in the brain datasets, EMB significantly outperforms SFL at every layer in Pereira2018 (0–24), NatStor_fMRI_ (0–24), and Tuckute2024 (1–24), and SFL never significantly outperforms EMB at any layer of any neural dataset, providing further support for the idea that brain responses reflect more direct access to rich, high-dimensional dynamics of language processing.

### **A.6** Dimensionality control

The two predictors we considered in this work have very different dimensionality: EMB has between 768 and 1,600 dimensions, whereas SFL has only three. It is thus possible that EMB might outperform SFL in the brain analyses not because it encodes richer linguistic content, but simply because a linear model over a high-dimensional feature space can express more complex response functions than a low-dimensional one—for instance, a nonlinear effect of surprisal. Note that this concern applies only to the neural results because in the behavioral datasets, SFL already outperforms EMB *despite* its smaller dimensionality.

To match the dimensionality of the two predictors, we projected SFL into a high-dimensional nonlinear feature space with random Fourier features (RFF; Rahimi and Recht, 2007). The three SFL predictors were mapped onto *D* random features, where *D* is the hidden size of the model under consideration, so that SFL_RFF_ has exactly the same dimensionality as EMB. We then repeated the analyses substituting SFL_RFF_ for SFL, keeping the rest of the procedure unchanged. For datasets where we had to pool estimates across words (thus, all but UCL_N400_), we obtained RFF representations at the word level and then summed them within each unit (sentence or TR).

The results are reported in Figure 6. EMB retained its advantage over SFL_RFF_ in four of the five neural datasets 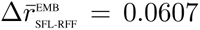, in line with our main findings, although in UCL_N400_ the two predictors obtained very similar performance. The only exception was Pereira2018, where SFL_RFF_ outperformed EMB (*−*0.0915). Importantly, even where SFL_RFF_ obtained strong results, EMB still accounted for substantial additional variance: concatenating EMB to SFL_RFF_ improved the model fit in all five neural datasets 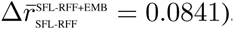, significantly so across all five models in UCL_N400_ (0.0796), Tuckute2024 (0.0889), and NatStor_fMRI_ (0.1406). The improvement was numerically similar but not statistically significant in Wehbe2014 (0.0816), the dataset with the lowest reliability in our sample (see Table 1), and both small and not significant in Pereira2018 (0.0297). Overall, the greater predictivity of EMB for brain responses cannot be reduced to its dimensionality as it predicts significant additional variance over a dimensionality-matched version of SFL.

**Figure 6:**
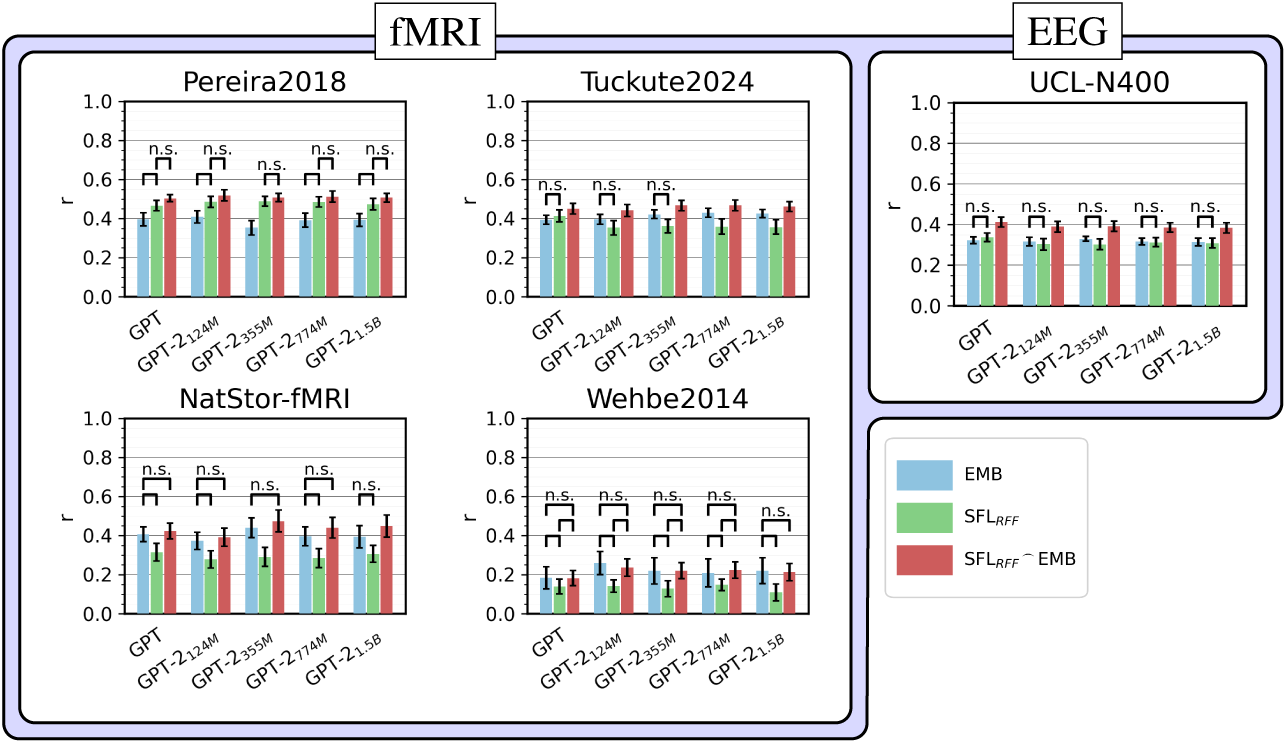
Predictivity of EMB, SFL_RFF_, and SFL_RFF⌢_EMB across the five neural datasets. SFL_RFF_ is a random Fourier feature expansion of surprisal, frequency, and length into the same number of dimensions as EMB. This expansion allows the effort-based predictors to express arbitrary nonlinear functions of SFL. EMB still outperforms SFL_RFF_ in four of the five neural datasets (though by a small margin in UCL_N400_). The concatenation SFL_RFF⌢_EMB improves over SFL_RFF_ alone in all datasets (though the difference was not significant in Pereira2018), indicating that the advantage of EMB over SFL for brain data is not an artifact of its dimensionality.

**Figure 7:**
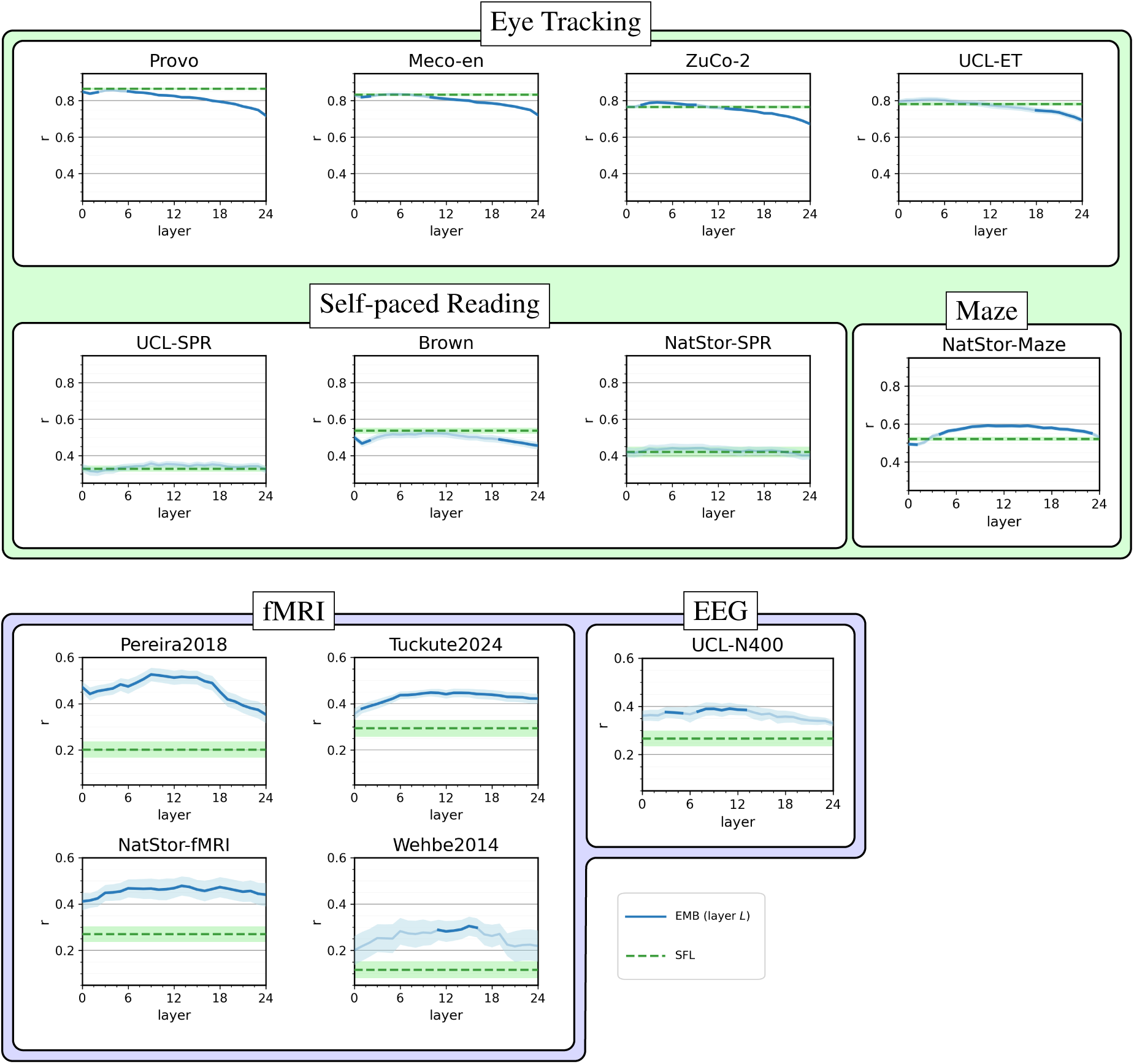
Predictivity of EMB extracted from each layer of GPT-2_355M_ (blue) compared to SFL (green dashed line). Shaded areas around the lines indicate the SEM over cross-validation folds. The blue line is faded where the difference between EMB and SFL is not statistically significant. In the eye-tracking datasets EMB obtains the strongest predictivity in early layers and then declines; in the self-paced reading datasets the two predictors obtain similar predictivity with the exception of Brown; in NatStor_Maze_ and all neural datasets EMB outperforms SFL.

A qualitative analysis of the model predictions further indicates that even if SFL_RFF_ and EMB obtain similar predictivity, they base their predictions on different features. The out-of-fold predictions of SFL_RFF_ and EMB are only moderately correlated with each other (Pereira2018: *r* = 0.39; UCL_N400_: *r* = 0.46; Tuckute2024: *r* = 0.56), and they encode different properties of the input. In Pereira2018—the dataset where SFL_RFF_ outperformed EMB—the predictions of SFL_RFF_ are strongly correlated with sentence length (*r* = 0.75), much more than those of EMB (*r* = 0.25); sentence length is itself a good predictor of the response in this dataset (*r* = 0.38). In Tuckute2024, the predictions of SFL_RFF_ mostly encode average word length (*r* = 0.42) and average word frequency (*r* = *−*0.36), whereas the predictions of EMB are less related to these surface properties (*r* = 0.19 and *r* = *−*0.12). Conversely, the predictions of EMB are more closely related to human ratings of imageability (*r* = *−*0.53) and of the extent to which the sentence describes something physical (*r* = *−*0.48) than those of SFL_RFF_ (*r* = *−*0.23 and *r* = *−*0.10). Thus, although a high-dimensional nonlinear expansion can increase the predictivity of SFL in some datasets (and decrease it in others), the resulting gains appear to be driven mostly by other shallow properties of the input, such as length and lexical frequency, whereas the predictions of EMB are more closely related to meaning.

### **A.7** Reliability analysis by sample size

To quantify the internal consistency of each dataset and assess how it scales with the number of observations, we computed progressive split-half reliability curves. For each dataset, we represented individual data points (using each dataset’s resolution, e.g., word-level reading times, TR-level fMRI responses, see Table 1) as lists of item-level measurements. We then iteratively estimated reliability for increasing numbers of observations (from 2 up to half the maximum available) using the following procedure. At each iteration, we randomly split the selected number of observations for each item into two halves, computed the average in each half, and correlated the resulting values across all items. This process was repeated 10 times to obtain stable estimates of reliability and associated standard deviations. The results are reported in Figure 8.

**Figure 8:**
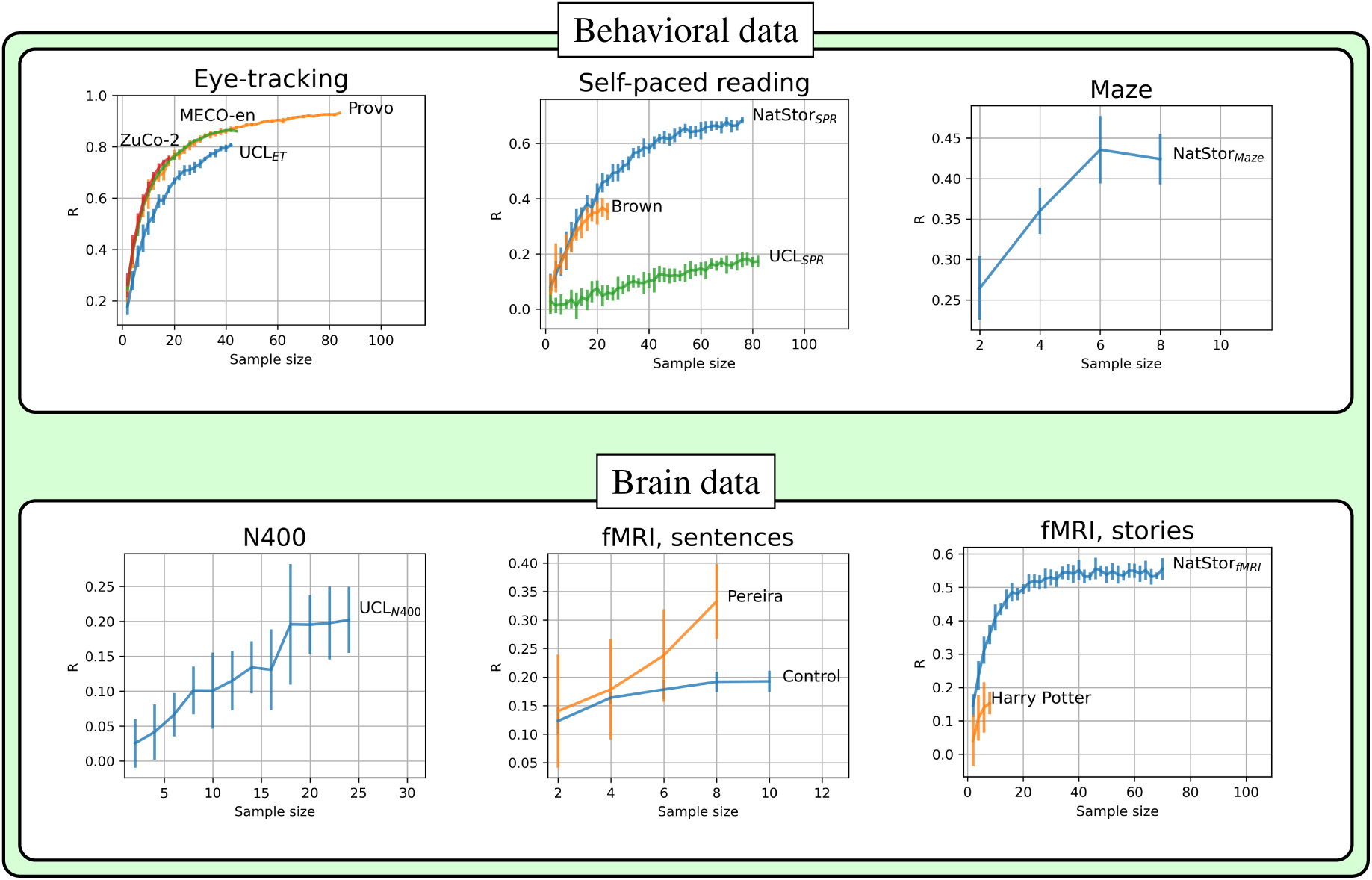
Progressive split-half reliability estimates across behavioral (top) and neural (bottom) datasets, as a function of sample size. For each dataset, reliability was calculated by randomly splitting item-level responses into two halves and correlating the means across groups. Plots are grouped by measurement type: eye-tracking, self-paced reading, maze, ERP (N400), and fMRI (sentence- or story-level designs).

^1^https://github.com/Andrea-de-Varda/SFL-EMB

